# UAV-based Remote Sensing of Bee Nesting Aggregations with Computer Vision for Object Detection

**DOI:** 10.1101/2025.06.16.659683

**Authors:** Tobias G. Mueller, Mark A. Buckner

## Abstract

1. Pollinating insects are in decline globally, threatening pollination services and driving a growing interest in pollinator monitoring and conservation. However, the implementation of conservation programs for these insects is often hindered by labor-intensive monitoring methods and in turn insufficient data to assess population trends.
2. We detail a method for surveying and censusing ground nesting bee aggregations, pairing automated UAV image capture with a custom trained computer vision-based object detection workflow using the YOLOv5m architecture. To highlight the ease of application and accuracy of the workflow, we surveyed a roughly 65m^2^ portion of a large *Colletes inaequalis* nesting aggregation. We compared the efficiency and performance of our model to manual counts of a technician.
3. Our model detected the location of 1,094 nests, representing 88% of the nests present in our test dataset, and a true positive rate of 97%. Adjusting for error, our model estimated a total of 1,250 nests across the study site, comparable to the total estimated from a manual count of 1,259 nests. Our model detected nests 20 times faster than the manual counts while mapping the aggregation with millimeter accuracy. Spatial analyses show that bee nest density was heterogenous, with dense spatially clustered regions comprised of upwards of 60 nests per m^2^.
4. *Synthesis and applications*: Our novel application of UAV imagery and object detection models for mapping and censusing a ground nesting bee aggregation represents a rapid, cost-effective solution for overcoming limitations in traditional manual methods. Our workflow generates essential data with the high throughput required to help inform the conservation decisions needed to stem global bee declines.

## Introduction

Human-caused biodiversity loss, fueled by multiple co-occurring drivers of global change, is contributing to the degradation of ecosystem services and faltering ecosystem function (Sage, 2020). Food security and global floral biodiversity rely on pollination services provided by animals, which pollinate approximately 90% of angiosperm species (Ollerton et al., 2011; Tong et al., 2023), with the majority pollination services provisioned by insects, especially bees (Willmer, 2011). In recent decades, rapid declines in bee populations and diversity have been linked to intensive pesticide use, habitat loss, invasive and pest species introductions, pathogens, and climate change (Dicks et al., 2021; Goulson et al., 2015; LeBuhn & Vargas Luna, 2021; Van Dooren, 2019; Zattara & Aizen, 2021). Concerns regarding declining bee diversity and the loss of bee-mediated pollination services has contributed to a growing interest in bee monitoring and conservation initiatives (Potts et al., 2024; Schlesinger et al., 2023; Woodard et al., 2020). Despite increasing research and policy centered on pollinator conservation, implementation of conservation programs for pollinating insects is often complicated by insufficient data on life histories, habitat preferences, physiology, and abundance.

Current studies of bee declines are often focused on determining trends in species diversity on large geographical scales (Bartomeus et al., 2013; Orr et al., 2021) and regional studies of local population are often performed with low spatial or temporal resolution, in part due to the effort required to carry out pollinator surveys. The current methods available for studying local bee populations are limited in their ability to discern population trends or environmental impacts on habitat quality. Presently, studies of bee populations often focus on foraging adults using structured sweep netting protocols (Brosi et al., 2008), bowl trapping (Kimoto et al., 2012), mark recapture (Hennessy et al., 2020), or visual observations (Larsson & Franzén, 2008). However, the emphasis on foraging adults does not necessarily capture the nesting population and may not accurately reflect population trends, with sampling results often influenced by local conditions at the time of sampling (Baum & Wallen, 2011; Packer & Darla-West, 2021). Removal methods and lethal sampling for identification may further confound estimates of population trends and potentially contribute to local declines, leading to a growing interest in alternative sampling methods (Miller et al., 2022; Montero-Castaño et al., 2022).

Research on ground nesting solitary bee aggregations is limited to a handful of traditional protocols. Research objectives focused on exploring nesting ecology are generally facilitated by manually marking and counting nests (Bischoff, 2003; Cameron et al., 1996; Linsley et al., 1952), quadrat studies (Dar et al., 2021; Giulian et al., 2024) or emergence traps (Sardiñas & Kremen, 2014). These methods, however, are laborious and time intensive and do not scale well to investigations of spatial or temporal variation. Additionally, the data collected are limited in their ability to account for fine grain variation in nest density and distributions and the resulting population estimates are coarse, often based on extrapolations from rough delineations of nest site extents, leading to observation error and uncertainty which could mask population trends. Despite the growing use of technology in pollinator research, the existing methods for investigating ground nesting bee nesting ecology at the site level remain largely unchanged. Advancing the study of nesting biology with fine spatial and temporal scale surveys is essential for informing bee conservation decisions and facilitating improvements in habitat restoration and conservation.

Novel technological innovations, such as automated remote sensing workflows, have the potential to move nesting surveys away from estimates and extrapolations, towards complete censuses of a site’s population at a fine spatial resolution. In comparison to traditional ground counts, manual counts of wildlife populations with unmanned aerial vehicle (UAV) imagery are more precise (Hodgson et al., 2016), however, manually counting of UAV imagery is time consuming and may give biased results (Torney et al., 2019). Deep learning object detection algorithms have become increasingly accessible with improvements in availability and abundance of ecological image data (Weinstein, 2018). Paired with the recent improvements and affordability of consumer level UAVs, there is a proliferation of studies exploring the application of automated UAV-based wildlife monitoring with object detection algorithms spanning a wide variety of objectives including surveys of ant mounds (dos Santos et al., 2022), sea bird nests counts (Cusick et al., 2024; Hayes et al., 2021), rare plant population censuses (Rominger & Meyer, 2021), and pest wasp monitoring (Jeong et al., 2023).

The introduction of inexpensive programmable UAVs with high resolution cameras, and the rapid advancement of artificial intelligence methods for object detection present an opportunity to increase the efficiency and coverage of ecological surveys of bee nesting aggregations. To test the feasibility and practicality of using automated UAV remote sensing to census wild bees, we surveyed a section of a large *Colletes inaequalis* nesting aggregation in Ithaca, New York, USA. Like 64% of described bee species, *C. inaequalis* is ground nesting (Cane & Neff, 2011), and has a propensity to form dense aggregations of hundreds to thousands of bees (López-Uribe et al., 2015). In the early spring, individuals emerge and mate, after which females tunnel into the ground, creating nests containing small underground chambers in which they provision pollen and lay eggs. Each nest belongs to a single active female and contains on average 3.5 eggs (Batra, 1980). From above, these nests appear as dark circular entrances surrounded by tumuli, piles of excavated dirt characteristic of many ground nesting species, creating a distinctive above ground signature.

To demonstrate the application of UAV imagery and computer vision-based nest detection, we detail a workflow for autonomous image acquisition and processing of an active *C. inaequalis* nesting aggregation. We apply a computationally tractable, open-source object detection modeling framework capable of identifying small objects across large orthomosaics. Here, we highlight the application of these methods for facilitating rapid, cost effective, and scalable censuses and millimeter scale spatial analyses to explore the population size and spatial structure of bee nesting aggregations with the throughput required to facilitate sub-daily repeat censuses. Finally, we discuss the application of UAV based surveys for ground nesting bee conservation and the avenues for future improvement of these methods. Our approach highlights the applicability of current, user-friendly object detection algorithms for modernizing the monitoring of known bee nesting aggregations and advancing our knowledge of bee life histories and nesting behavior.

## Methods

We imaged the *C. inaequalis* nesting aggregation on 23 April 2024, during active nest provisioning, with an Air2S UAV (SZ DJI Technology Co., Ltd.), a consumer grade drone with a 20MP, 1-inch CMOS camera sensor. To facilitate autonomous flight and image capture, we generated a series of waypoints with the Flight Planner plugin (Gruca, 2023) in QGIS (QGIS Development Team, 2024). We placed route waypoints with 70% image overlap covering a large extent of the nesting aggregation that was representative of the variability in nest and vegetation density. The routing was set to follow terrain at a constant elevation of two meters above ground level (AGL) based on a one meter resolution digital elevation model (New York Office of Information Technology Services, 2021), resulting in a nominal ground sampling distance of 0.6mm/pixel. We programmed an autonomous flight plan from the waypoint coordinates (WGS84) with Litchi (VC Technology Ltd, 2024). At each waypoint, we set a two second delay allowing the UAV to stabilize prior to image capture with a gimbal angle of −90 degrees (pointing directly towards the ground). All images were merged into a single orthomosaic using WebODM v2.5.2 (Toffanin et al., 2024).

On the same day, we used the UAV to capture images across the nesting aggregation, but outside the surveyed area of interest, to develop a representative training dataset. To ensure the model was robust to changes in altitude, lighting, and vegetation, we captured images that represented the variation in vegetation and lighting conditions at altitudes between one and three meters AGL. Training images were split into 608×608 pixel images for annotation, resulting in a comprehensive training dataset of 1,512 images that included a range of nest densities and background images. We labeled all active nests within the training set using Label Studio (Tkachenko et al., 2020) and we included nests at the edge of the image only when the entrance hole was clearly identifiable and not cut off.

We implemented our bee nest detection workflow with YOLOv5, a pretrained object detection model using the "you only look once" model architecture (Redmon et al., 2016) under the Ultralytics framework (Jocher et al., 2022). We employed the standard architecture for the medium model, which we trained to detect nest entrances using our training image set with 300 epochs with batch gradient descent. To compensate for the small size of the nests within the image we employed Slicing Aided Hyper Inference (SAHI) (Akyon et al., 2022). SAHI is a generic framework which can be combined with any other object detection algorithms to improve the detection of small objects by slicing images into a series of overlapping patches with increased relative pixel sizes. The resulting patches are fed through the object detection algorithm’s forward pass before the bounding boxes are merged across overlapping patches to return all detections across the original input image. We sliced the image at 608×608 pixels, matching the training image size, with a 40% overlap between slices.

To assess model performance, we split the orthomosaic along with the model detections into 608×608 pixel tiles and randomly selected 40% of tiles (365 images) which we manually labeled in Label Studio to act as a test set (Figure S1). In evaluating the model against the test set, we employed a workflow using Voxel51 (Moore & Corso, 2020). We defined true positives as model prediction bounding boxes that had an intersection over union (IOU) of 0.4, i.e., the proportion of overlap between the prediction bounding box and the ground truth bounding box. Since nests have ill-defined boundaries and are often spatially isolated from one another, a higher IOU is not required for differentiating true positives from false positives and the use of a higher IOU may lead to some true positive detections being incorrectly labeled as false positives. We calculated the confidence level which maximized F1 score (the harmonic mean between precision and recall) and evaluated model performance with F1, precision, recall, and mean average precision (mAP) at this optimal confidence level. All nest detections across the full orthomosaic were thresholded at the determined optimal confidence level, dropping any detections with a confidence score below the lower limit. To assess our model’s robustness to image acquisitions at higher altitudes, and therefore larger ground sample distances and lower resolutions, we downsampled the orthomosaic to half the resolution, approximating a flight of twice the altitude, and performed the same workflow as described above.

To facilitate further analysis, we drew on the ability of the object detections model to localize the predicted nest bounding boxes. We first georeferenced the orthomosaic using the GPS coordinates of five ground control points spread throughout the study area and projected the image to WGS84 / UTM zone 18N (EPSG:32618). To estimate the point locations for each nest detection, we calculated the centroids of each individual bounding box. We examined the density and distribution of the nests throughout the study area with spatstat v3.3.1 (Baddeley et al., 2015) in R v4.4.3 (R Core Team, 2025). We mapped the density of nests with kernel density estimation using a bandwidth selected to minimize the mean-square error criterion (Berman & Diggle, 1989). To identify if the distribution of nests differs from complete spatial randomness across the study area, we calculated the local L-function, a local linearized version of Ripley’s K-function also called the neighborhood density function, at a distance of 0.1m (Baddeley et al., 2015; Getis & Franklin, 1987). Finally, we calculated the nearest neighbor distance as the Euclidean distance from each point to its nearest neighbor, similar to the data collected in the field during a previous quadrat-based study of a nesting aggregation’s distribution pattern (Potts & Willmer, 1998).

To evaluate how our detection model compared to manual methods, we calculated the difference in accuracy and time costs with manual nest counts performed by an undergraduate technician. The technician was provided training on nest identification, counting methods, and given examples of edge cases, including emergence holes and abandoned nests. After training, the technician was instructed to count all nests across the full orthomosaic using DotDotGoose v1.7.0 (Ersts, 2024) and record the time required to complete the task. To assess the performance of the manual counts, the resulting point locations of all counted nest locations were manually transformed into bounding boxes and compared against the labeled test set images. For comparison with the object detection model, we calculated the F1 score, precision, recall, and mAP of the manual counts as described above.

## Results

The orthomosaic from the UAV imagery covered a 65.3m^2^ (5.1m by 12.8m) section of the *C. inaequalis* nesting aggregation, representing substantial variation in nest and vegetation density. Our YOLOv5 and SAHI object detection workflow detected 1,094 nests across the study area. Based on our model’s predictive performance across the test dataset, we can estimate the true count within the study area is ∼1,250 nests. Assuming an average of 3.5 brood cells per nest, we estimate a total of 4,375 individuals in the next generation prior to overwintering mortality.

The time to import the 791.7 MB orthomosaic, run the detection model, and export the predictions was 2 minutes and 38 seconds when run on a GPU (1152 CUDA cores; 3GB) and 7 minutes and 52 seconds when run on a CPU (6 core; 3.6 GHz). Both tests were run using consumer grade components and 64GB DDR4-3200 RAM. The resultant model predictions had an F1 score of 0.921 (precision 0.969; recall 0.877) when thresholded at the optimal confidence level of 0.625. The model mAP was 0.457. This translates to 97% of the model nest predictions being true nests and the model finding 88% of the nests present in the test dataset. Overall, the model predictions included few false positives, however, the model slightly underestimated the total number of nests present in the study area.

False positives were generally caused by dirt-colored leaves of similar size to the nest tumuli (Figure 1a) as well as mounds of dirt perhaps caused by other organisms or old unused nests (Figure 1b) that did not have adequate representation in the training images. False negatives often occurred when two nests were directly adjacent to one another and their tumuli greatly overlapped with the model only detecting one of the nests (Figure 1d). The model sometimes grouped both overlapping nests into a single large detection resulting in the detection being classified as two false negatives and as a false positive during model evaluation (Figure 1c). The model also occasionally failed to detect nests that were partially covered by debris or vegetation (Figure 1e) or surrounded by large patches of dirt without a distinct tumulus (Figure 1f).

**Figure 1.**
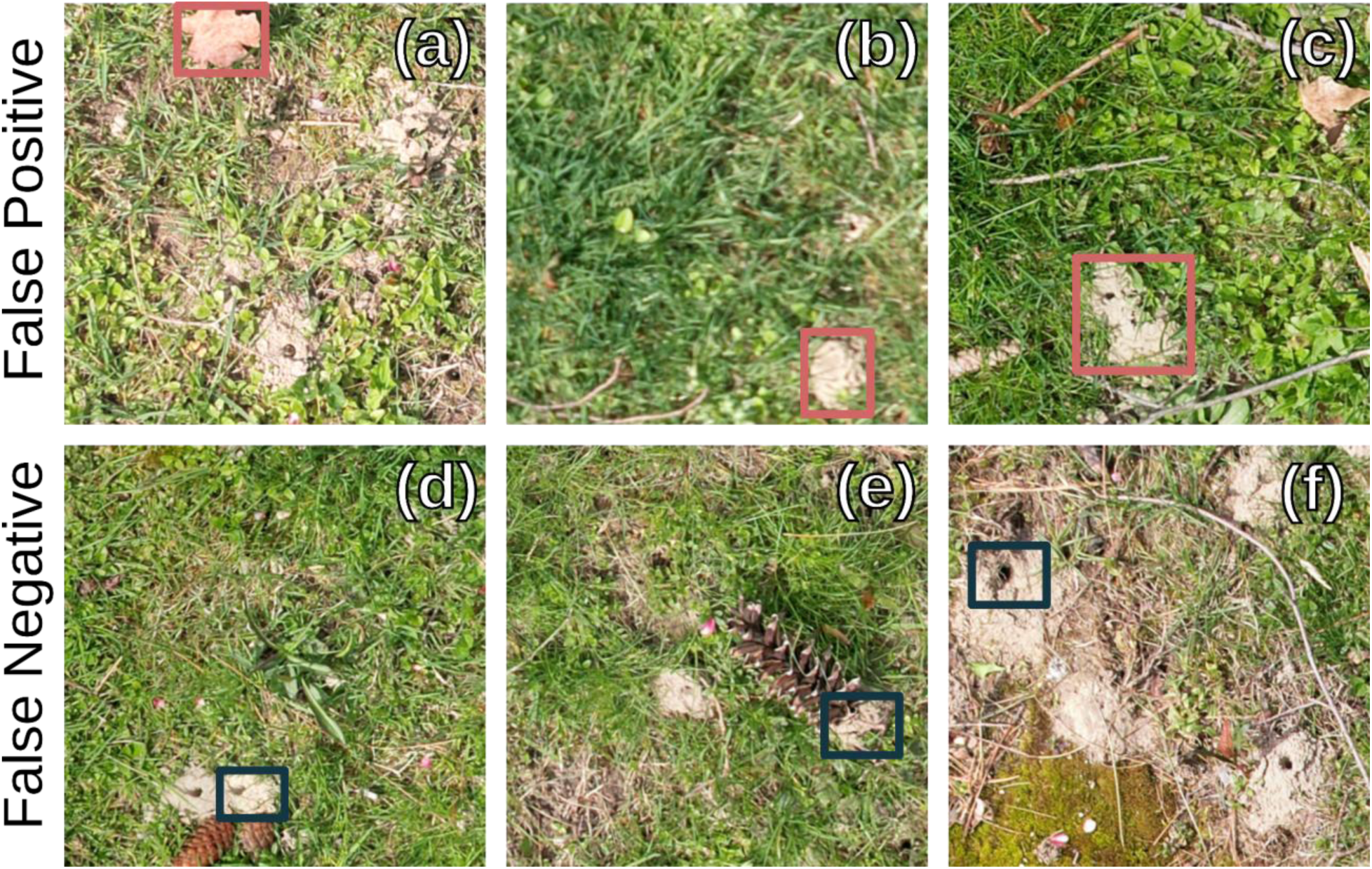
False positives (3% of predictions) were caused by (a) unusual leaves sized similarly to nests; (b) old nests that were no longer active; (c) two overlapping nests detected as one combined nest. False negatives (12% of ground truth nests) were also caused by (d) overlapping nests; (e) nests partially covered by grass or debris; (f) nests in a larger patch of dirt with low or nonexistent tumuli. Overlapping nests sometimes led to both a false positive and a false negative detection.

The model was able to detect nests in images from down-sampled imagery representing the expected resolution from a flight at double the altitude, with a decrease in run time of 76% to 37 seconds. Despite the lower resolution and the mismatch between training and test image resolution, the model performed comparably well when predicting across the low-resolution dataset with an F1 of 0.903 (precision = 0.97, recall = 0.85) at an optimal confidence level of 0.263.

In evaluating our model against manual counts by a trained undergraduate technician, we found comparable model performance. The object detection model out performed manual counts with fewer false positive detections contributing to an approximate 8 percentage point increase in precision relative to the manual counts (manual: 0.89, model: 0.97) and a comparable, through slightly lower recall (manual: 0.90, model: 0.88). In total, the undergraduate technician counted 1,259 nests, consistent with our adjusted model estimate of 1,250 nests. The principal difference between these two approaches was in the throughput and total task completion time. The undergraduate technician annotated and counted the full orthomosaic in just under an hour, 51.5 minutes, representing a throughput of 131.4 hours per hectare. The model processed the full resolution orthomosaic in 2.6 minutes using an average consumer-grade computer, almost 20 times faster, representing a theoretical throughput of 6.7 hr/ha, assuming linear scaling. With the low-resolution dataset, the task completion time was 84 times faster, albeit with the tradeoff of reduced recall compared to the manual counts.

The kernel density map, estimated with a bandwidth of 0.38 m, showed two distinct regions of high-density nesting (Figure 2a). Nest density peaked at approximately 62 nests/m^2^ in the central high-density nesting zone and a lower density of 53 nests/m^2^ in the second most dense region. The Local L-function calculated with a distance of 0.1m, indicated nest clustering (Figure 2b) in the densest regions. Throughout the remainder of the study area the Local L-functions suggested most nests were clustered, but to a lesser degree, with values above the expectation for complete spatial randomness. This pattern held true across distances down to below 2.5 cm (Figure S2). Within these dense clusters, most nests were within 10 cm of their nearest neighbor with distances increasing towards the outside of the defined clusters and towards the western edge of the aggregation (Figure 2c).

**Figure 2.**
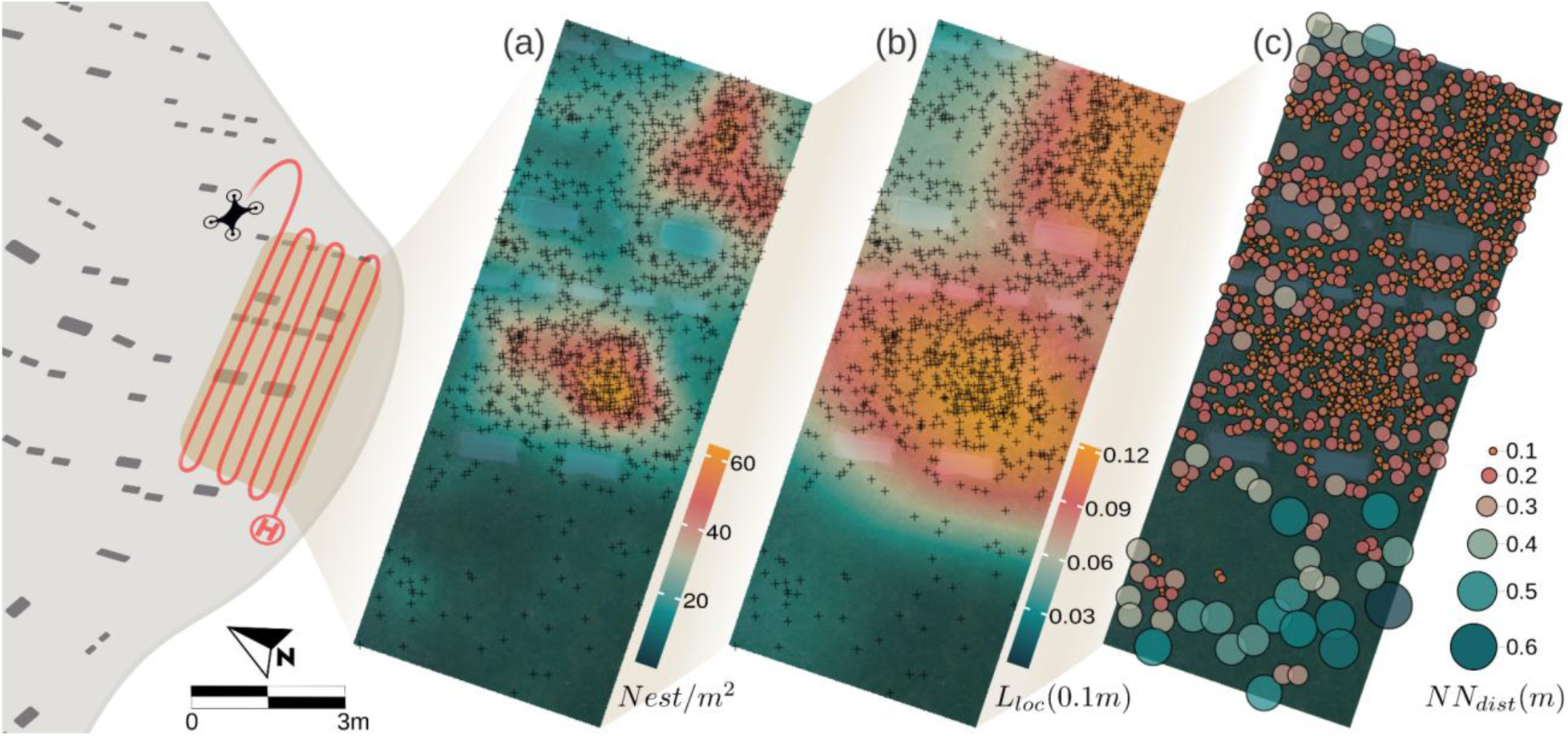
Spatial point pattern analyses of the *C. inaequalis* nesting aggregation within the study area. (a) Nest density, calculated with kernel density estimation, revealed two distinct high-density patches of the study area. (b) Local L-function indicated that the nests are more clustered than expected under complete spatial randomness at 0.1m, with clustering decreasing towards the edge of the nesting aggregation in correspondence with decreasing density. (c) Nearest neighbor distances, represented as a Steinen diagram, demonstrate that the majority of nests within the highly clustered regions were within 20cm of the next closest nest and 10 cm or less in the densest areas.

## Discussion

Traditional methods for researching ground nesting bee aggregations are laborious and do not scale well to broad temporal or spatial studies, particularly in large aggregations which may be comprised of millions of nests (Blagoveshchenskaya, 1963). Here, we present a rapid, cost effective, and precise method based on an open-source object detection model and autonomous UAV remote sensing to map a large bee nesting aggregation with millimeter scale accuracy. In benchmarking the nest detection model against manual counts, we showed that UAV-based mapping can greatly reduce processing time by upwards of 2,000%, opening the door to repeat daily, or sub-daily censuses of nest site use, nesting behavior, and comprehensive spatial analyses of large aggregations. Our model-based methods performed comparably to manual counts and were better able to resolve true detections, with the tradeoff of a minor underestimation in the proportion of nests identified. The model results show that it tends to undercount the total number of nests, resulting in conservative estimates of population sizes. In the context of population monitoring for at-risk species, overestimation may lead to inaction or delayed action and a bias towards underestimating population counts may be preferable to overestimations. The model had a very high precision of 0.97, meaning almost all model detections are true nests. This is critical for population surveys where low precision could contribute to erroneous abundance estimates and over confidence in population stability. Further refinement of model structure and training could address the slightly lower recall resulting from the model’s inability to detect some cryptic nests, presenting a potential focal area for future improvement.

This model was trained solely to detect nests of a single bee species, *Colletes inaequalis*, and was not designed to differentiate between, or detect across, nests of different species. However, many bees exhibit a similar nesting morphology, with a simple dirt mound created when excavating a central tunnel and the model may perform well across many bee species which share this morphology. Using the same described methodology, the existing model can easily and iteratively be trained to detect nests across species or even to differentiate among species. This may be particularly feasible for distinguishing between bees with distinct above-ground nest structures, such as some species in the genus *Diadasia*, which construct tall chimney-like structures in place of the more common unstructured tumuli (Eickwort et al., 1977).

Remote detection of bee nests may not be feasible for all of the ∼64% of bee species which excavate nests (Cane & Neff, 2011). Some species are extremely cryptic, nesting under thick vegetation or rocks (Jackson et al., 2025) or sealing their nest entrances with dirt or debris between visits (Hurd et al., 1974). While these cases represent a potential limitation of our approach, many ground nesting species are thought to prefer open grassy or sandy areas with little tall vegetation, suggesting that this methodology can be applied to a substantial proportion of bee species (Antoine & Forrest, 2021; Potts et al., 2005). However, nesting biology alone may not be the only limiting factor, weather can impact the external presentation of nests, making them more difficult to detect which may be particularly important for studies of species which nest during the rainy season. For example, following heavy rains in the spring in New York the tumuli of *C. inaequalis* nests can be temporarily washed away removing the above ground signature the model was trained to recognize. We also note that remote sensing does not eliminate the need for traditional life history investigations and nest excavations, particularly for understanding below ground nest structure, reproductive rates, and brood cell provisioning, among others. Our novel surveying methods, however, work in conjunction with natural history observations to make analyses of bee nesting aggregations more repeatable and cost effective.

Additionally, the YOLO family of models, and other general object detection models, may not perform optimally with many small and heavily clustered objects (Akyon et al., 2022; Redmon et al., 2016; Yang et al., 2019). We mediated this issue effectively by predicting across the study area with SAHI, but did note false negatives from neighboring nests with overlapping tumuli, an issue which may become exacerbated in particularly dense nesting aggregations. A larger more diverse suit of training images may be able to better capture the variation in nest structure, aid in the differentiation of nests from their surrounding substrate, and improve the model’s ability to discriminate between bee nests and other objects resembling tumuli, e.g., anthills, worm castings, leaves, etc. Alternatively, other object detection models architectures may be more suitable for the unique task of nest identification and may provide improved performance over YOLOv5. However, the application of other model architectures may be demanding in comparison, and the use of the Ultralytics framework, as we employed here, is a popular and user-friendly implementation of object detection for ecological problems, enabling the adoption of this method by researchers without extensive experience in computer vision.

While there are opportunities to refine the model architecture to optimize bee nest detection and generalize to a greater diversity of bee nest morphologies, this study successfully demonstrates the efficiency of using UAVs and object detection to census large bee nesting aggregations with high accuracy. While we only selected a subset of the aggregation as our area of interest, the use of an automated flight plan allows for the easy scaling of this workflow to cover significant areas with minimal additional effort. With our model proving robust to changes in image resolution, we suggest that the altitude at which images are captured can be increased by at least double, significantly reducing the time required for image acquisition, the overall dataset size, and further improving computational efficiency. Moreover, the application of real-time object detection using video sequences from a UAV presents an opportunity to more efficiently scale-up remote nest detection. Such an implementation would simplify flight planning and may reduce the time needed for image acquisition. Animal detection in UAV videos is already being implemented in other ecological systems (Van Gemert et al., 2015) and likely could be adapted for bee nest identification.

Given the ongoing concerns related to pollinator conservation, innovative technology like UAV-based nest detection can accelerate targeted conservation and research efforts for ground nesting bees. The method we present here enables population censusing at extremely fine spatial and temporal scales, a level of detail that current techniques cannot tractably achieve. The flexibility of this method enables studies of a broad array of bee species within their natural habitat, including those which are not easily managed or lab reared. This opens the possibility to evaluate habitat quality and within-site use based on localized variation in soils, pesticides, vegetation, or microclimatic factors. Deconstructing how bees interact with microhabitats will enable granular risk assessments and inform local restoration and protection initiatives through revealing fine-scale environmental preferences and enabling risk mitigation for known disturbance events, e.g. construction. The rapid and repeatable nature of our automated data acquisition and processing workflow enables efficient data collection across time, capturing intra- and interdiel and -annual variation in how threats impact individual populations and behavior. UAV-based monitoring of aggregations across a disturbance event, for instance a period of extreme heat or land-use change, can facilitate in situ assessments of nesting activity disruption. Additionally, since the data is spatially explicit, it would allow for the identification of spatial variation of risk within a nesting aggregation and could guide habitat restoration for species of conservation concern.

Our workflow pairing automated UAV imaging with a custom trained object detection model provides a rapid and cost-effective solution for censusing ground nesting bee aggregations. We show significant improvements in spatial resolution, and reduction in effort over traditional manual methods without sacrificing accuracy, highlighting that this affordable workflow, utilizing consumer level technology, is a promising method for generating meaningful conservation data. The limitations of current monitoring approaches have hindered bee conservation action necessitating a modernization of methods (Portman et al., 2020). By embracing conservation technology and computational methods, UAV-based remote sensing of bee nesting aggregations provides a novel approach for direct monitoring of bee population declines.

## Acknowledgements

We would like to thank Cherie and Kevin Morse, managers of the East Lawn Cemetery in Ithaca, NY, for allowing us to study the nesting aggregation located on their property. Additionally, we would like to thank Carrie Day for logistical support, Daniel Sorokin for his time manually counting bee nests, as well as Bryan Danforth, Heather Grab, and Kevin Li for their feedback on the manuscript. This work was supported by the Cornell Atkinson Center for Sustainability’s Graduate Research Grant. TGM was funded through NSF GRFP #DGE – 2139899.

## Data Availability Statement

The object detection model training dataset and processed orthomosaic are archived and openly available on Figshare (Mueller & Buckner, 2025a) along with all associated code and the trained nest detection model (Mueller & Buckner, 2025b) which is also available on GitHub (https://github.com/bcknr/AIggregation).

## Author Contributions

Tobias G. Mueller: Conceptualization, Data curation, Formal Analysis, Funding acquisition, Investigation, Methodology, Resources, Software, Visualization, Writing – original draft, Writing – review & editing; Mark A. Buckner: Conceptualization, Data curation, Formal Analysis, Investigation, Methodology, Resources, Software, Visualization, Writing – original draft, Writing – review & editing

## References

Akyon, F. C., Onur Altinuc, S., & Temizel, A. (2022). Slicing Aided Hyper Inference and Fine-Tuning for Small Object Detection. 2022 IEEE International Conference on Image Processing (ICIP), 966–970. 10.1109/ICIP46576.2022.9897990

Antoine, C. M., & Forrest, J. R. K. (2021). Nesting habitat of ground-nesting bees: A review. Ecological Entomology, 46(2), 143–159. 10.1111/een.12986

Baddeley, A., Rubak, E., & Turner, R. (2015). Spatial Point Patterns: Methodology and Applications with R. Chapman and Hall/CRC Press. https://www.routledge.com/Spatial-Point-Patterns-Methodology-and-Applications-with-R/Baddeley-Rubak-Turner/p/book/9781482210200/

Bartomeus, I., Ascher, J. S., Gibbs, J., Danforth, B. N., Wagner, D. L., Hedtke, S. M., & Winfree, R. (2013). Historical changes in northeastern US bee pollinators related to shared ecological traits. Proceedings of the National Academy of Sciences of the United States of America, 110(12), 4656–4660. 10.1073/pnas.1218503110

Batra, S. W. T. (1980). Ecology, Behavior, Pheromones, Parasites and Management of the Sympatric Vernal Bees Colletes inaequalis, C. thoracicus and C. validus. Journal of the Kansas Entomological Society, 53(3), 509–538.

Baum, K. A., & Wallen, K. E. (2011). Potential Bias in Pan Trapping as a Function of Floral Abundance. Journal of the Kansas Entomological Society, 84(2), 155–159. 10.2317/JKES100629.1

Berman, M., & Diggle, P. (1989). Estimating Weighted Integrals of the Second-Order Intensity of a Spatial Point Process. Journal of the Royal Statistical Society: Series B (Methodological), 51(1), 81–92. 10.1111/j.2517-6161.1989.tb01750.x

Bischoff, I. (2003). Population dynamics of the solitary digger bee Andrena vaga Panzer (Hymenoptera, Andrenidae) studied using mark-recapture and nest counts. Population Ecology, 45(3), 197–204. 10.1007/s10144-003-0156-6

Blagoveshchenskaya, N. (1963). Giant Colony of the Solitary Bee Dasypoda Plumipes PZ. (Hymenoptera, Melittidae). Entomological Review, 42(1), 60–61.

Brosi, B. J., Daily, G. C., Shih, T. M., Oviedo, F., & Durán, G. (2008). The effects of forest fragmentation on bee communities in tropical countryside. Journal of Applied Ecology, 45(3), 773–783. 10.1111/j.1365-2664.2007.01412.x

Cameron, S. A., Whitfield, J. B., Hulslander, C. L., Cresko, W. A., Isenberg, S. B., & King, R. W. (1996). Nesting Biology and Foraging Patterns of the Solitary Bee Melissodes rustica (Hymenoptera: Apidae) in Northwest Arkansas. Journal of the Kansas Entomological Society, 69(4), 260–273.

Cane, J. H., & Neff, J. L. (2011). Predicted fates of ground-nesting bees in soil heated by wildfire: Thermal tolerances of life stages and a survey of nesting depths. Biological Conservation, 144(11), 2631–2636. 10.1016/j.biocon.2011.07.019

Cusick, A., Fudala, K., Storożenko, P. P., Świeżewski, J., Kaleta, J., Oosthuizen, W. C., Pfeifer, C., & Bialik, R. J. (2024). Using machine learning to count Antarctic shag (*Leucocarbo bransfieldensis*) nests on images captured by remotely piloted aircraft systems. Ecological Informatics, 82, 102707. 10.1016/j.ecoinf.2024.102707

Dar, S. A., Sofi, M. A., El-Sharnouby, M., Hassan, M., Rashid, R., Mir, S. H., Naggar, Y. A., Salah, M., Gajger, I. T., & Sayed, S. (2021). Nesting behaviour and foraging characteristics of *Andrena cineraria* (Hymenoptera: Andrenidae). Saudi Journal of Biological Sciences, 28(8), 4147–4154. 10.1016/j.sjbs.2021.04.063

Dicks, L. V., Breeze, T. D., Ngo, H. T., Senapathi, D., An, J., Aizen, M. A., Basu, P., Buchori, D., Galetto, L., Garibaldi, L. A., Gemmill-Herren, B., Howlett, B. G., Imperatriz-Fonseca, V. L., Johnson, S. D., Kovács-Hostyánszki, A., Kwon, Y. J., Lattorff, H. M. G., Lungharwo, T., Seymour, C. L.,… Potts, S. G. (2021). A global-scale expert assessment of drivers and risks associated with pollinator decline. Nature Ecology & Evolution, 5(10), 1453– 1461. 10.1038/s41559-021-01534-9

dos Santos, A., Biesseck, B. J. G., Latte, N., de Lima Santos, I. C., dos Santos, W. P., Zanetti, R., & Zanuncio, J. C. (2022). Remote detection and measurement of leaf-cutting ant nests using deep learning and an unmanned aerial vehicle. Computers and Electronics in Agriculture, 198, 107071. 10.1016/j.compag.2022.107071

Eickwort, G. C., Eickwort, K. R., & Linsley, E. G. (1977). Observations on Nest Aggregations of the Bees Diadasia olivacea and D. diminuta (Hymenoptera: Anthophoridae). Journal of the Kansas Entomological Society, 50(1), 1–17.

Ersts, P., J. (2024). DotDotGoose (Version 1.7.0) [Computer software]. American Museum of Natural History, Center for Biodiversity and Conservation. https://biodiversityinformatics.amnh.org/open_source/dotdotgoose

Getis, A., & Franklin, J. (1987). Second-Order Neighborhood Analysis of Mapped Point Patterns. Ecology, 68(3), 473–477. 10.2307/1938452

Giulian, J., Danforth, B. N., & Kueneman, J. G. (2024). A Large Aggregation of Melissodes bimaculatus (Hymenoptera: Apidae) Offers Perspectives on Gregarious Nesting and Pollination Services. Northeastern Naturalist, 31(3), 402–417. 10.1656/045.031.0314

Goulson, D., Nicholls, E., Botías, C., & Rotheray, E. L. (2015). Bee declines driven by combined stress from parasites, pesticides, and lack of flowers. Science, 347(6229), 1255957. 10.1126/science.1255957

Gruca, J. (2023). Flight Planner (Version 0.6.1) [Computer software]. https://github.com/JMG30/flight_planner

Hayes, M. C., Gray, P. C., Harris, G., Sedgwick, W. C., Crawford, V. D., Chazal, N., Crofts, S., & Johnston, D. W. (2021). Drones and deep learning produce accurate and efficient monitoring of large-scale seabird colonies. Ornithological Applications, 123(3), duab022. 10.1093/ornithapp/duab022

Hennessy, G., Goulson, D., & Ratnieks, F. L. W. (2020). Population assessment and foraging ecology of nest aggregations of the rare solitary bee, Eucera longicornis at Gatwick Airport, and implications for their management. Journal of Insect Conservation, 24(6), 947–960. 10.1007/s10841-020-00266-8

Hodgson, J. C., Baylis, S. M., Mott, R., Herrod, A., & Clarke, R. H. (2016). Precision wildlife monitoring using unmanned aerial vehicles. Scientific Reports, 6(1), 22574. 10.1038/srep22574

Hurd, P. D., Linsley, E. G., & Michelbacher, A. D. (1974). Ecology of the squash and gourd bee, Peponapis pruinosa, on cultivated cucurbits in California (Hymenoptera: Apoidea). Smithsonian Contributions to Zoology, 168, 1–17. 10.5479/si.00810282.168

Jackson, F. M., Prendergast, K. S., Hardy, G., & Xu, W. (2025). Enhancing Lasioglossum (Homalictus) dotatum (Hymenoptera: Halictidae) habitats: The role of rock gravel in bare soil landscapes. Austral Entomology, 64(2), e70008. 10.1111/aen.70008

Jeong, Y., Jeon, M.-S., Lee, J., Yu, S.-H., Kim, S., Kim, D., Kim, K.-C., Lee, S., Lee, C.-W., & Choi, I. (2023). Development of a Real-Time Vespa velutina Nest Detection and Notification System Using Artificial Intelligence in Drones. Drones, 7(10), Article 10. 10.3390/drones7100630

Jocher, G., Chaurasia, A., Stoken, A., Borovec, J., NanoCode012, Kwon, Y., Michael, K., TaoXie, Fang, J., imyhxy, Lorna, Yifu), 曾逸夫(Zeng, Wong, C., V, A., Montes, D., Wang, Z., Fati, C., Nadar, J., Laughing,… Jain, M. (2022). ultralytics/yolov5: V7.0 - YOLOv5 SOTA Realtime Instance Segmentation (Version v7.0) [Computer software]. Zenodo. 10.5281/zenodo.7347926

Kimoto, C., DeBano, S. J., Thorp, R. W., Rao, S., & Stephen, W. P. (2012). Investigating temporal patterns of a native bee community in a remnant North American bunchgrass prairie using blue vane traps. Journal of Insect Science, 12(1), 108. 10.1673/031.012.10801

Larsson, M., & Franzén, M. (2008). Estimating the population size of specialised solitary bees. Ecological Entomology, 33(2), 232–238. 10.1111/j.1365-2311.2007.00956.x

LeBuhn, G., & Vargas Luna, J. (2021). Pollinator decline: What do we know about the drivers of solitary bee declines? Current Opinion in Insect Science, 46, 106–111. 10.1016/j.cois.2021.05.004

Linsley, E. G., MacSwain, J. W., & Smith, R. F. (1952). Outline for Ecological Life Histories of Solitary and Semi-Social Bees. Ecology, 33(4), 558–567. 10.2307/1931531

López-Uribe, M. M., Morreale, S. J., Santiago, C. K., & Danforth, B. N. (2015). Nest Suitability, Fine-Scale Population Structure and Male-Mediated Dispersal of a Solitary Ground Nesting Bee in an Urban Landscape. PLOS ONE, 10(5), e0125719. 10.1371/journal.pone.0125719

Miller, Z. J., Lynn, A., Oster, C., Piotter, E., Wallace, M., Sullivan, L. L., & Galen, C. (2022). Unintended Consequences? Lethal Specimen Collection Accelerates with Conservation Concern. American Entomologist, 68(3), 48–55. 10.1093/ae/tmac057

Montero-Castaño, A., Koch, J. B. U., Lindsay, T.-T. T., Love, B., Mola, J. M., Newman, K., & Sharkey, J. K. (2022). Pursuing best practices for minimizing wild bee captures to support biological research. Conservation Science and Practice, 4(7), e12734. 10.1111/csp2.12734

Moore, B. E., & Corso, J. J. (2020). FiftyOne. GitHub. Note: https://Github.Com/Voxel51/Fiftyone.

Mueller, T. G., & Buckner, M. A. (2025a). UAV-based Remote Sensing of Bee Nesting Aggregations with Computer Vision for Object Detection [Dataset]. figshare. 10.6084/m9.figshare.29308523

Mueller, T. G., & Buckner, M. A. (2025b). UAV-based Remote Sensing of Bee Nesting Aggregations with Computer Vision for Object Detection [Computer software]. 10.6084/m9.figshare.29309684

New York Office of Information Technology Services. (2021, March). Lidar Collection (QL2) of all or part of Schuyler, Seneca, Steuben, Tompkins, Wayne and Yates Counties, NY Lidar; Hydro-Flattened Bare-Earth DEM. https://gis.ny.gov/nys-dem

Ollerton, J., Winfree, R., & Tarrant, S. (2011). How many flowering plants are pollinated by animals? Oikos, 120(3), 321–326. 10.1111/j.1600-0706.2010.18644.x

Orr, M. C., Hughes, A. C., Chesters, D., Pickering, J., Zhu, C.-D., & Ascher, J. S. (2021). Global Patterns and Drivers of Bee Distribution. Current Biology, 31(3), 451–458.e4. 10.1016/j.cub.2020.10.053

Packer, L., & Darla-West, G. (2021). Bees: How and Why to Sample Them. In J. C. Santos & G. W. Fernandes (Eds.), Measuring Arthropod Biodiversity: A Handbook of Sampling Methods (pp. 55–83). Springer International Publishing. 10.1007/978-3-030-53226-0_3

Portman, Z. M., Bruninga-Socolar, B., & Cariveau, D. P. (2020). The State of Bee Monitoring in the United States: A Call to Refocus Away From Bowl Traps and Towards More Effective Methods. Annals of the Entomological Society of America, 113(5), 337–342. 10.1093/aesa/saaa010

Potts, S. G., Bartomeus, I., Biesmeijer, K., Breeze, T., Casino, A., Dauber, J., Dieker, P., Hochkirch, A., Høye, T., Isaac, N., Kleijn, D., Laikre, L., Mandelik, Y., Montagna, M., Montero, C. A., Öckinger, E., Oteman, B., Pardo, V. A., Polce, C.,… Zhang, J. (2024). Refined proposal for an EU pollinator monitoring scheme. JRC Publications Repository. 10.2760/2005545

Potts, S. G., Vulliamy, B., Roberts, S., O’Toole, C., Dafni, A., Ne’eman, G., & Willmer, P. (2005). Role of nesting resources in organising diverse bee communities in a Mediterranean landscape. Ecological Entomology, 30(1), 78–85. 10.1111/j.0307-6946.2005.00662.x

Potts, S. G., & Willmer, Pat. (1998). Compact housing in built-up areas: Spatial patterning of nests in aggregations of a ground-nesting bee. Ecological Entomology, 23(4), 427–432. 10.1046/j.1365-2311.1998.00160.x

QGIS Development Team. (2024). QGIS Geographic Information System. QGIS Association. https://www.qgis.org

R Core Team. (2025). R: A Language and Environment for Statistical Computing. R Foundation for Statistical Computing. https://www.R-project.org/

Redmon, J., Divvala, S., Girshick, R., & Farhadi, A. (2016). You Only Look Once: Unified, Real-Time Object Detection. 2016 IEEE Conference on Computer Vision and Pattern Recognition (CVPR), 779–788. 10.1109/CVPR.2016.91

Rominger, K. R., & Meyer, S. E. (2021). Drones, Deep Learning, and Endangered Plants: A Method for Population-Level Census Using Image Analysis. Drones, 5(4), Article 4. 10.3390/drones5040126

Sage, R. F. (2020). Global change biology: A primer. Global Change Biology, 26(1), 3–30. 10.1111/gcb.14893

Sardiñas, H. S., & Kremen, C. (2014). Evaluating nesting microhabitat for ground-nesting bees using emergence traps. Basic and Applied Ecology, 15(2), 161–168. 10.1016/j.baae.2014.02.004

Schlesinger, M. D., White, E. L., Corser, J. D., Danforth, B. N., Fierke, M. K., Greenwood, C. M., Hatfield, R. G., Hietala-Henschell, K. G., Mawdsley, J. R., McFarland, K. P., Niver, R., Rozen, J. G., Van Dyke, M., & Howard, T. G. (2023). A multi-taxonomic survey to determine the conservation status of native pollinators. Frontiers in Ecology and Evolution, 11. 10.3389/fevo.2023.1274680

Tkachenko, M., Malyuk, M., Holmanyuk, A., & Liubimov, N. (2020). Label Studio: Data labeling software. https://github.com/HumanSignal/label-studio

Toffanin, P., Leo, L. D., Chamo, N., Farell, K., Saijin-Naib, Barker, B., Mather, S., Joseph, D., bot, pyup io, Kaluza, O., Davis, B., IZem, A., Bateman, C., Islam, T., Poulain, S., USPLM, Acuña, D., Machado, R. W., Ves, N.,… Heggy, I. (2024). OpenDroneMap/WebODM: 2.5.0 (Version v2.5.0) [Computer software]. Zenodo. 10.5281/zenodo.11193946

Tong, Z.-Y., Wu, L.-Y., Feng, H.-H., Zhang, M., Armbruster, W. S., Renner, S. S., & Huang, S.-Q. (2023). New calculations indicate that 90% of flowering plant species are animal-pollinated. National Science Review, 10(10), nwad219. 10.1093/nsr/nwad219

Torney, C. J., Lloyd-Jones, D. J., Chevallier, M., Moyer, D. C., Maliti, H. T., Mwita, M., Kohi, E. M., & Hopcraft, G. C. (2019). A comparison of deep learning and citizen science techniques for counting wildlife in aerial survey images. Methods in Ecology and Evolution, 10(6), 779–787. 10.1111/2041-210X.13165

Van Dooren, T. J. M. (2019). Assessing species richness trends: Declines of bees and bumblebees in the Netherlands since 1945. Ecology and Evolution, 9(23), 13056–13068. 10.1002/ece3.5717

Van Gemert, J. C., Verschoor, C. R., Mettes, P., Epema, K., Koh, L. P., & Wich, S. (2015). Nature Conservation Drones for Automatic Localization and Counting of Animals. In L. Agapito, M. M. Bronstein, & C. Rother (Eds.), Computer Vision—ECCV 2014 Workshops (Vol. 8925, pp. 255–270). Springer International Publishing. 10.1007/978-3-319-16178-5_17

VC Technology Ltd. (2024). Litchi for DJI Drones. https://apps.apple.com/us/app/litchi-for-dji-drones/id1059218666

Weinstein, B. G. (2018). A computer vision for animal ecology. Journal of Animal Ecology, 87(3), 533–545. 10.1111/1365-2656.12780

Willmer, P. (2011). Pollination and Floral Ecology. Princeton University Press. 10.1515/9781400838943

Woodard, S. H., Federman, S., James, R. R., Danforth, B. N., Griswold, T. L., Inouye, D., McFrederick, Q. S., Morandin, L., Paul, D. L., Sellers, E., Strange, J. P., Vaughan, M., Williams, N. M., Branstetter, M. G., Burns, C. T., Cane, J., Cariveau, A. B., Cariveau, D. P., Childers, A.,… Wehling, W. (2020). Towards a U.S. national program for monitoring native bees. Biological Conservation, 252, 108821. 10.1016/j.biocon.2020.108821

Yang, F., Fan, H., Chu, P., Blasch, E., & Ling, H. (2019). Clustered Object Detection in Aerial Images (No. arXiv:1904.08008). arXiv. 10.48550/arXiv.1904.08008

Zattara, E. E., & Aizen, M. A. (2021). Worldwide occurrence records suggest a global decline in bee species richness. One Earth, 4(1), 114–123. 10.1016/j.oneear.2020.12.005

