## Supplementary material for "UAV-based Remote Sensing of Bee Nesting Aggregations with Computer Vision for Object Detection": figure S1

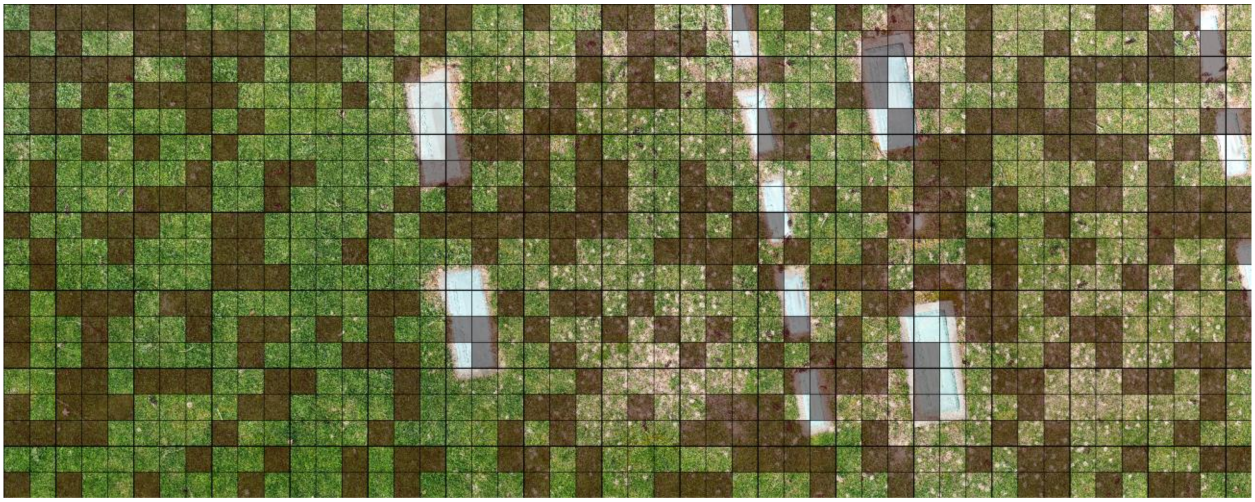

**Figure S1-** A test set consisting of 40% of the study site, 365 tiles shown in red, was randomly selected to evaluate model performance.

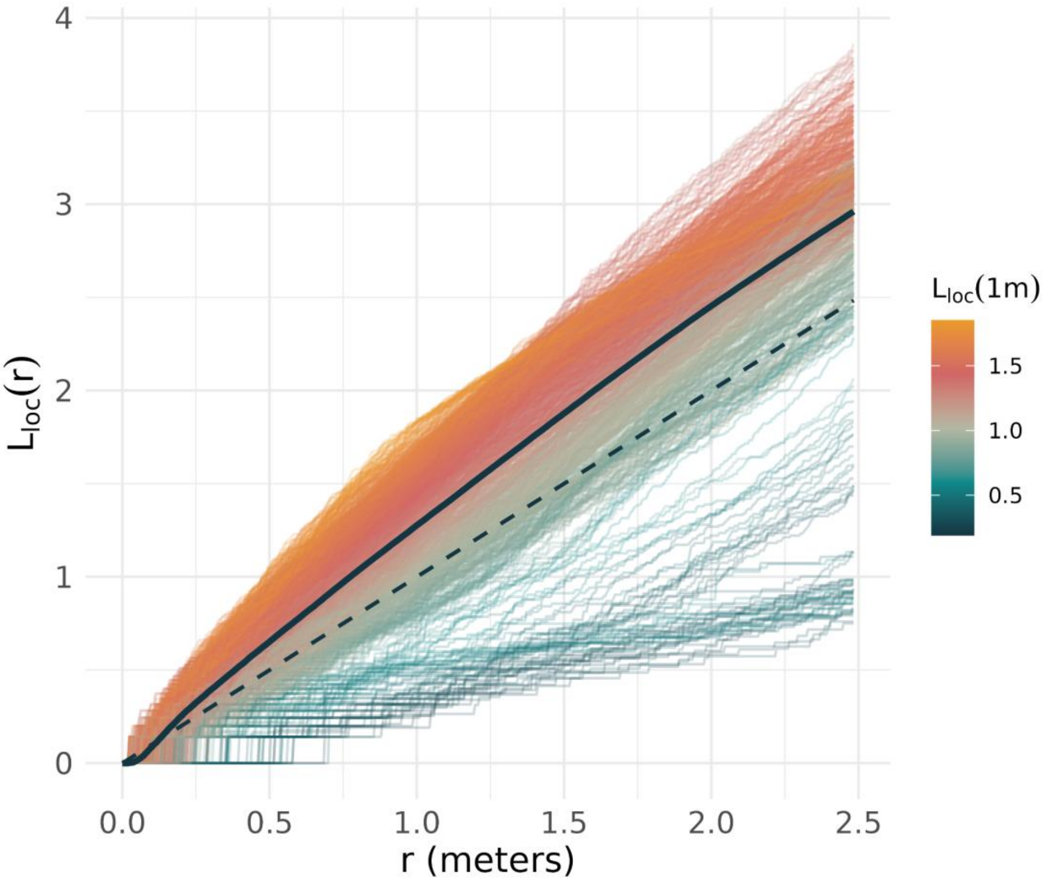


**Figure S2-** Plot of local L-functions for all nests at distances (r) up to 2.5m with line color corresponding to the value of the function when r = 1 m. Lines below the dashed theoretical line are dispersed and lines above are clustered. The distance which a line remains along the x-axis ($L_{loc}(r) = 0$) is the nearest neighbor distance, showing that the nearest neighbor distance increases with more dispersed nests. The mean value (solid blue line) of all nests in the study area is more clustered than expected under complete spatial randomness with the average nest showing clustering ($L_{loc}(r) > r$) above approximately 2.5cm.
